# Comment on ‘Accumbens cholinergic interneurons dynamically promote dopamine release and enable motivation’

**DOI:** 10.1101/2023.12.27.573485

**Authors:** James Taniguchi, Riccardo Melani, Lynne Chantranupong, Michelle J. Wen, Ali Mohebi, Joshua Berke, Bernardo Sabatini, Nicolas Tritsch

**Affiliations:** Neuroscience Institute and Fresco Institute for Parkinson’s and Movement Disorders, New York University Grossman School of Medicine, New York, USA; Department of Neurobiology, Howard Hughes Medical Institute, Harvard Medical School, Boston, USA; Department of Neurology, University of California, San Francisco, San Francisco, USA

## Abstract

Acetylcholine is widely believed to modulate the release of dopamine in the striatum of mammals. Experiments in brain slices clearly show that synchronous activation of striatal cholinergic interneurons is sufficient to drive dopamine release via axo-axonal stimulation of nicotinic acetylcholine receptors. However, evidence for this mechanism *in vivo* has been less forthcoming. A recent paper in *eLife* (Mohebi *et al*., 2023) reported that, in awake behaving rats, optogenetic activation of striatal cholinergic interneurons with blue light readily evokes dopamine release measured with the red fluorescent sensor RdLight1. Here, we show that blue light alone alters the fluorescent properties of RdLight1 in a manner that may be misconstrued as phasic dopamine release, and that this artefactual photoactivation can account for the effects attributed to cholinergic interneurons. Our findings indicate that measurements of dopamine using the red-shifted fluorescent sensor RdLight1 should be interpreted with caution when combined with optogenetics. In light of this and other publications that did not observe large acetylcholine-evoked dopamine transients *in vivo*, the conditions under which such release occurs in behaving animals remain unknown.

## Introduction

Presynaptic modulation is a ubiquitous mechanism through which neural circuits control the amount of chemical transmitter that axons release per incoming action potential (Lovinger et al., 2022). Striatum-projecting midbrain dopamine (DA) neurons are no exception; many transmitters present in the striatum directly act on DA axons to facilitate or depress vesicular release of DA (Sulzer et al., 2016). However, the axons of DA neurons do stand out in their sensitivity to acetylcholine (ACh), whose release from striatal cholinergic interneurons in brain slices is potent enough to trigger axonal action potentials and locally evoke DA release (i.e., independently of somatic spiking activity) via the activation of β2-containing nicotinic ACh receptors on DA axons (Cachope et al., 2012; Threlfell et al., 2012; Wang et al., 2014; Mamaligas et al., 2016; Kramer et al., 2022; Liu et al., 2022; Matityahu et al., 2023). This mechanism was proposed to underlie observations that DA release in ventral striatum (nucleus accumbens, NAc) can increase even when the firing of DA neurons in the midbrain appears unchanged (Berke 2018; Mohebi et al., 2019). However, direct evidence for ACh-evoked striatal DA release *in vivo* remains limited.

In their original report of ACh-evoked DA release, Cachope and colleagues (2012) presented an example recording whereby strong optogenetic activation of cholinergic interneurons for several seconds is accompanied by DA elevation in the NAc of a urethane-anesthetized mouse. More recently, we and others showed in mice and rats that the patterns of DA and ACh release in various striatal locations *in vivo* are strongly correlated on sub-second time-scales in a manner that is consistent with ACh-evoked DA release (Howe et al., 2019; Liu et al., 2022; Chantranupong et al., 2023; Krok et al., 2023; Mohebi et al., 2023). Yet DA dynamics in the dorsal and lateral striatum were found to persist even after molecular, pharmacological and optogenetic interference with ACh signaling (Chantranupong et al., 2023; Krok et al., 2023).

A recent paper in *eLife* (Mohebi et al., 2023) provided some of the most compelling evidence to date that optogenetic activation of channelrhodopsin-expressing striatal cholinergic interneurons drives DA release in the NAc of awake behaving rats, as measured with the red-shifted DA sensor RdLight1 (Patriarchi et al., 2020). However, one concern with these experiments is that mApple-based fluorescent sensors – including RdLight1 and the GRAB-rDA3 series (Zhuo et al., 2023) – may exhibit photoactivation (also known as ‘photoswitching’ or ‘photoconversion’), a process whereby mApple’s red fluorescence changes in the presence of blue light (Shaner et al., 2008). This phenomenon is one of the main downsides of the R-GECO family of red Ca^2+^ indicators, which also use mApple and grow brighter independently of Ca^2+^ for hundreds of milliseconds following brief flashes of blue light, limiting their use with optogenetics (Akerboom et al., 2013; Dana et al., 2016). In the case of RdLight1, it was previously shown that photoactivation effects are negligible when expressed in cultured kidney cells and imaged using light-scanning confocal microscopy (Patriarchi et al., 2020; but see Zhuo et al., 2023). Whether RdLight1 shows photoactivation *in vivo* under conditions routinely used in behavioral experiments (i.e. fiber photometry and full-field optogenetic stimulation) has not been investigated.

## Results

To determine if blue light modifies the fluorescent properties of RdLight1 in the behaving brain, we virally expressed RdLight1 in either the dorsolateral striatum (DLS; N=8) or NAc (N=8) of wild-type mice (**Figure 1A,B**) and imaged RdLight1 fluorescence *in vivo* using fiber photometry while mice were head-fixed on a cylindrical treadmill (**Figure 1C**). Under standard continuous illumination conditions (565 nm excitation light, 30-50 μW at the tip of the fiber), we observed transient increases and decreases in red fluorescence consistent with established patterns of DA release (Howe and Dombeck 2016; Da Silva et al., 2018; Wei et al., 2022; Chantranupong et al., 2023; Krok et al., 2023; Markowitz et al., 2023), including spontaneous fluctuations during immobility, movement-related transients during self-paced wheel running and large-amplitude reward-evoked responses (not shown).

**Figure 1.**
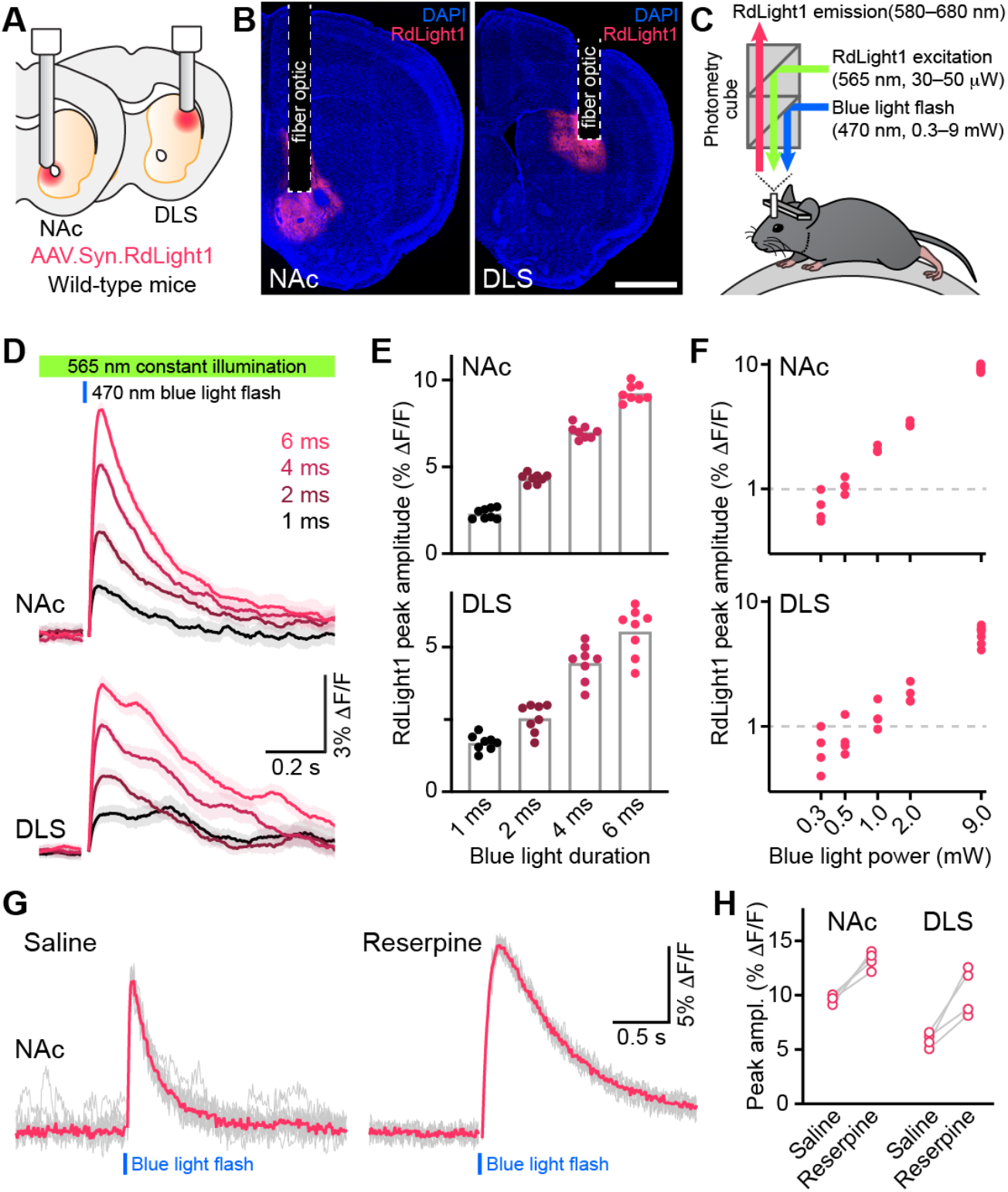
Blue light evokes RdLight1 photoactivation transients resembling DA release. **(A)** The red DA sensor RdLight1 was virally expressed in neurons of the nucleus accumbens (NAc) or dorsolateral striatum (DLS) of wild-type mice and imaged *in vivo* via chronically-implanted fiber optic cannulas. **(B)** Example fixed coronal sections from two mice stained for the nuclear marker DAPI (blue) and imaged by epifluorescence showing RdLight1 (red) expression in the NAc (left) and DLS (right). Scale bar: 1 mm. **(C)** Experimental setup for imaging RdLight1 by photometry while delivering blue light (470 nm) via the same fiber optic in awake behaving mice. RdLight1 was excited with continuous yellow-green light (565 nm) and red emitted fluorescence (580–680 nm) was collected via a dual color photometry minicube. **(D)** RdLight1 fluorescence (emitted upon continuous excitation with 565 nm light) increases upon exposure to blue light pulses (470 nm; 9 mW at tip of patch cord) of various durations (1–6 ms; color-coded) in the NAc (top) and DLS (bottom) of a representative mouse. Solid lines are the mean of 10 blue light presentations. Shaded area shows SEM. **(E)** The magnitude of blue light-evoked RdLight1 fluorescence transients (i.e., photoactivation) grows with the duration of blue light pulses in both NAc (top; N = 8 mice) and DLS (bottom; N = 8 mice).**(F)** Magnitude of RdLight1 fluorescence transients evoked by 6 ms-long blue light pulses of various intensities (0.3–9 mW, measured at tip of patch cord) in both NAc (top) and DLS (bottom). Data plotted on Log_10_-Log_10_ scales. **(G)** Representative RdLight1 photoactivation transients (red: mean of 10 individual traces shown in gray) imaged in NAc before and after systemic block of vesicular DA release with reserpine. **(H)** Magnitude of blue light-evoked (9mW; 6 ms pulse width) RdLight1 photoactivation before and after reserpine treatment in NAc (N = 4 mice) and DLS (N = 4 mice).

Delivering blue light pulses through the same fiber at powers typically used for optogenetic manipulations (6 ms-long; 9 mW at the tip of the patch cord) evoked distinct transients in RdLight1 fluorescence resembling DA release (**Figure 1D**). In the NAc, these transients averaged 9.3 ± 0.2 % ΔF/F in magnitude, peaked 112 ± 2 ms after light onset and decayed back to baseline within 1 s (τ_decay_: 438 ± 24 ms; N=8 mice). In the DLS, blue light-evoked transients were smaller (5.6 ± 0.3 % ΔF/F) but showed similar kinetics (time from light onset to peak: 135 ± 5 ms; τ_decay_: 467 ± 43 ms; N=8 mice). In both regions, blue light-evoked transients scaled in magnitude with the duration (**Figure 1E**) and intensity of light pulses (**Figure 1F**). In a separate laboratory, we observed comparable increases in RdLight1 fluorescence in N=3 mice that expressed RdLight1 in the ventrolateral striatum (VLS; **Figure 1 – Figure Supplement 1**), confirming their occurrence across a range of experimental conditions. These delayed fluorescent signals are specific to RdLight1, as they are not observed in mice expressing the red fluorescent protein tdTomato (not shown).

Do these blue light-evoked RdLight1 transients reflect a phasic elevation in extracellular DA? Several lines of evidence suggest that this is not the case. First, our wild-type mice do not express blue light-gated opsins to drive DA release and DA neurons are not thought to be intrinsically sensitive to blue light. Second, blue light-evoked transients show little trial-by-trial variability in their amplitudes and kinetics (**Figure 1D,G**). Third, blue light-evoked transients do not display short-term facilitation or depression under a variety of stimulation conditions (**Figure 2** and **Figure 1 – Figure Supplement 1C**), calling into question their synaptic origin. Fourth, we repeated the above experiments in a subset of mice treated with reserpine, an irreversible antagonist of the transporter required for loading DA into synaptic vesicles (**Figure 1G**). Under these conditions, spontaneous fluctuations in RdLight1 fluorescence vanished in both the DLS (N=4) and NAc (N=4), confirming the absence of activity-dependent DA release *in vivo*. By contrast, blue light-evoked RdLight1 transients did not disappear and, if anything, grew in amplitude and duration in both DLS and NAc (**Figure 1G,H**), demonstrating that they do not reflect synaptic release of DA.

**Figure 2.**
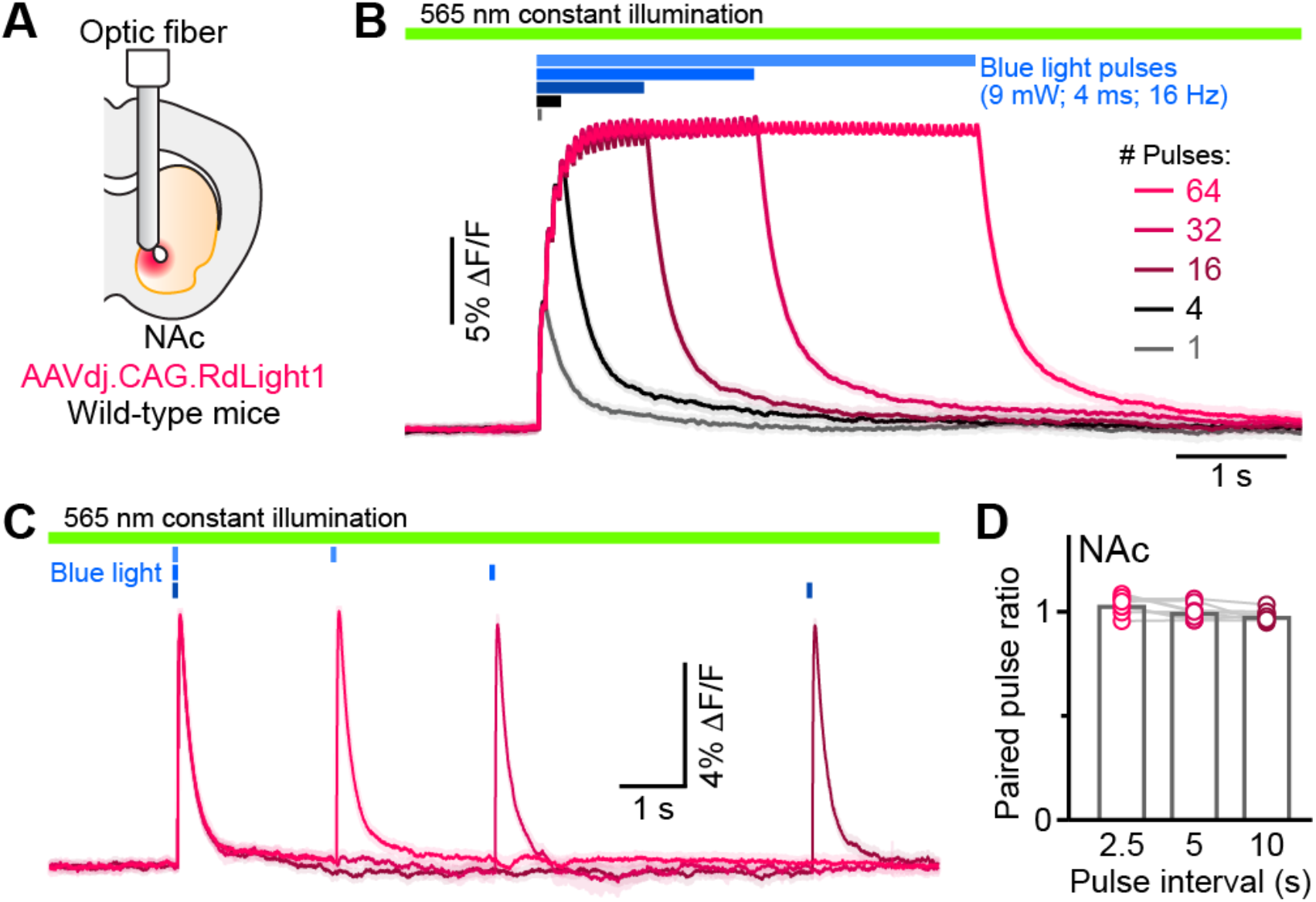
RdLight1 photoactivation transients evoked by blue light pulse trains. **(A)** Experimental setup. **(B)** Mean (± SEM) RdLight1 photoactivation in response to 1, 4, 16, 32, or 64 blue light pulses (9 mW, 4 ms width, 16 Hz frequency) in the NAc of a representative mouse. Green bar illustrates constant 565 nm illumination. Blue bars show when blue light is delivered. **(C)** Same as (B) for pairs of blue light pulses (9 mW, 4 ms width) separated by 2.5, 5 and 10 s.**(D)** Mean paired-pulse ratio (magnitude of pulse #2/pulse #1) across three inter-pulse intervals in each of N=8 mice.

## Discussion

Collectively, our results show that RdLight1 displays strong photoactivation following exposure to blue light under conditions routinely used to monitor and manipulate neural activity *in vivo*. This photoactivation manifests as a prolonged, DA-independent increase in RdLight1 fluorescence that outlasts the blue light pulse and slowly decays back to baseline over hundreds of milliseconds, giving it the appearance of synaptically-released DA. Under our recording conditions, photoactivation remained detectable with as little as 0.3 mW blue light (**Figure 1F**), indicating that RdLight1 fluorescence should be interpreted with caution when combined with blue light in a variety of experimental conditions, including dual-color imaging with alternating stimulation of green and red fluorophores.

In addition, our findings call into question the nature of the RdLight1 fluorescent transients reported in Figure 1 of the study by Mohebi et al. (2023). Given the similarity of our recordings in terms of response magnitude, timing and dynamics over a variety of stimulation parameters, it is likely that the light-evoked RdLight1 responses reported reflect this photoactivation effect. Although the study used 405 nm illumination to control for changes in fluorophore properties independent of ligand binding (i.e., the so-called isosbestic point), this deep blue wavelength has not been shown to be the isosbestic point for red-shifted fluorophores such as RdLight1. Additional experiments will therefore be needed to determine the conditions under which cholinergic interneurons locally evoke DA release in the striatum of behaving animals. If using RdLight1, experiments should include a mutated sensor that does not bind DA, controls expressing RdLight1 only (i.e., no opsin, to control for the effects of blue light alone) and minimal constant blue light stimulation. Alternatively, green fluorescent protein (GFP)-based DA sensors such as dLight1 (Patriarchi et al., 2018) or GRAB-DA3 (Zhuo et al., 2023) may be used in combination with red-shifted optogenetic actuators (Klapoetke et al., 2014; Marshel et al., 2019), although this configuration is not without caveats either, as blue light illumination may cause opsin activation.

## Materials and Methods

### Animals

Procedures were performed in accordance with protocols approved by the NYU Grossman School of Medicine (NYUGSM) and Harvard Medical School (HMS) Institutional Animal Care and Use Committees. Wild type mice (C57Bl6/J; Jackson Laboratory strain #000664; 12–18 weeks of age) were housed in group before surgery and singly after surgery under a reverse 12-hour light-dark cycle (dark from 6 a.m. to 6 p.m. at NYUGSM and 10 a.m. to 10 p.m. at HMS) with *ad libitum* access to food and water.

### Stereotaxic surgery

Mice were prepared for intracranial infections of adeno-associated viruses (AAVs) as before (Chantranupong et al., 2023; Krok et al., 2023). Briefly, mice were anaesthetized with isoflurane, administered Ketoprofen (10 mg/kg, subcutaneous) or Carprofen (5 mg/kg, subcutaneous) and placed in a stereotaxic apparatus (Kopf Instruments), where a small craniotomy was drilled above the NAc (from Bregma: AP +1.0 mm, ML +0.75 mm), DLS (from Bregma: AP +0.5 mm, ML +2.5 mm), or VLS (from Bregma: AP +0.6 mm, ML +2.3 mm). 300 nL of AAV2/9.Syn.RdLight1 (CNP Viral Vector Core at the CERVO Research Center contribution) was injected (100 nL/min) at a depth of 3.7 mm below dura for NAc, 2.3 mm for DLS or 3.2 mm for VLS using a microsyringe pump (KD Scientific; Legato 111) connected to a pulled glass injection needle (100 μm tip; Drummond Wiretrol II). Fiber optics (NAc and DLS: 400 μm diameter core, 0.5 NA; RWD Life Science Inc; VLS: 200 μm diameter core, 0.48 NA; Doric) were implanted 100 μm above the injection site and cemented to the skull using C&B metabond (Parkell) along with a custom titanium headpost placed over lambda to allow for head fixation during recordings. Mice were allowed to recover in their home cage for 2–4 weeks before head-fixation and treadmill habituation, and recordings.

### Fiber Photometry

RdLight1 photometry recordings were carried out by feeding constant, low-power yellow-green excitation light (565 nm LED, 30–50 μW at the tip of the patch cord; Thorlabs M565F3) to a fluorescence mini cube (FMC5_E1(460-490)_F1(500-540)_E2(555-570)_F2(580-680)_S; Doric) connected to the mouse’s fiber optic implant via a 0.48 NA patch cord (NAc and DLS: MFP_400/460/900-0.48_2m_FCM-MF1.25; VLS: MFP_200/220/900_2m_FCM-MF1.25; both from Doric). The red light emitted by RdLight1 was collected through the same patch cord and routed via the fluorescence mini cube and a second fiber optic (MFP_600/630/LWMJ-0.48_0.5m_FCM-FCM; Doric) to a photoreceiver (Newport 2151) to produce a voltage that is proportional to the intensity of the emitted light. Voltages were digitized at 2 kHz with either a National Instruments acquisition board (NI USB-6343) or a Labjack (T7) and saved to disk as ‘trials/sweeps’ lasting 5–20 s in duration each using Wavesurfer sotware (Janelia). To characterize the photoactivation behavior of RdLight1, we delivered brief pulses (1–6 ms in duration) of blue light (NAc, DLS: 470 nm LED, Thorlabs M470F3; VLS: 470 nm laser, Optoengine) to the brain via the same fluorescence mini cube and patch cord used for photometry. We tested a range of blue light powers (measured at the tip of the patch cord) frequently used for optogenetic manipulations *in vivo*. The timing, duration and intensity of blue light pulses were controlled digitally using Wavesurfer, with each stimulation parameter repeated at minimum 10 times per mouse/recording site. RdLight1 photoactivation responses are extremely stable over time and were reliably seen for the duration of 1 h-long recording sessions.

### Fiber Photometry Analysis

Photometry signals were processed and analyzed offline using custom code in MATLAB (Mathworks) and Igor Pro 6.02A (Wavemetrics). Raw voltages collected from the photoreceiver were down sampled to 20 Hz and converted to ‘percent changes in fluorescence’ using the equation 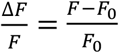, where *F*_*0*_ is the mean baseline fluorescence, computed for each trial/sweep during a 1.5–2 s baseline window preceding the blue light stimulus. Blue excitation light led to an instantaneous artifact in the red channel for the exact duration of the blue light pulse, which was blanked in display panels. For each experiment, 10–12 replicates were performed. In figures, gray traces represent single trials, while colored traces represent the mean of 10–12 trials, with standard error of the mean (SEM) shown as a shaded area. The properties of RdLight1 photoactivation transients [peak amplitude, latency to peak (i.e., from blue light onset to RdLight1 photoactivation peak) and decay time constant (i.e., time from peak to 37% of peak)] were measured for each mouse using averaged waveforms. Data are reported in the text and figures as mean ± SEM. N-values represent the number of mice.

### Immunohistochemistry

Mice were deeply anesthetized with isoflurane and perfused transcardially with 4% paraformaldehyde in phosphate buffered saline (PBS). Brains were post-fixed for 24 h and sectioned coronally (100 μm in thickness) using a vibratome (Leica; VT1000S). Brain sections were mounted on superfrost slides and coverslipped with ProLong antifade reagent with DAPI (Molecular Probes). RdLight1 fluorescence was not immuno-enhanced. Whole sections were imaged with an Olympus VS120 slide scanning microscope.

### Reagents

To inhibit DA vesicular transport and prevent vesicular release of DA, mice were injected intraperitoneally with the irreversible vesicular monoamine transporter inhibitor reserpine (5 mg/kg) for 24 h prior to RdLight1 photometry.

## Acknowledgments

This work was supported by the National Institutes of Health (R01MH130658 to N.X.T), a National Science Foundation (NSF) Graduate Research Fellowship (to J.T) and a Hirschl-Weill-Caulier Career Scientist Award (to N.X.T). We acknowledge the New York University Langone Health Department of Comparative Medicine for animal care and maintenance and the Neuroscience Institute’s imaging facilities for microscope availability.

## Author contributions

B.L.S. and N.X.T conceived of the project. J.T. and R.M. performed experiments in the NAc and DLS. L.C. and M.J.W. carried out experiments in the VLS. J.T., R.M., L.C., B.L.S and N.X.T. analyzed and interpreted the data, and wrote the manuscript in collaboration with A.M and J.B.

---

**Figure 1 – Figure supplement 1.**
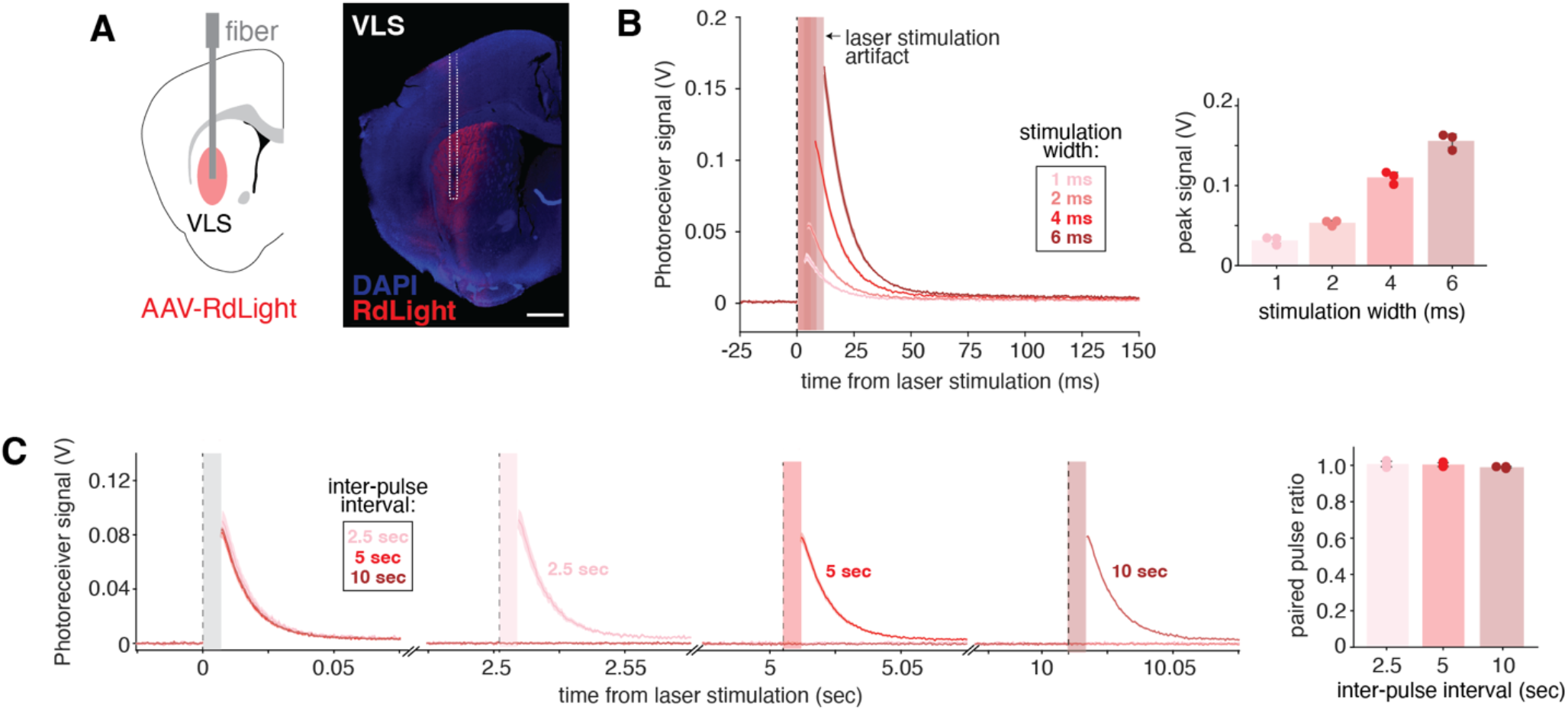
Blue light evokes RdLight1 photoactivation in VLS. **(A)** Injection and fiber implantation scheme for mice expressing RdLight1 in the VLS (let). Images of sensor expression from a representative mouse (right; scale bar: 800 μm). **(B)** Mean (± SEM) RdLight1 photoactivation signal in the VLS in response to 1, 2, 4, or 6 ms single pulses of blue light stimulation (10 mW) coupled with constant 565 nm illumination (30 μW). Colored vertical bars indicate laser stimulation artifacts which were removed. Mean maximal RdLight1 photoactivation signals recorded under the indicated stimulation widths are displayed. Each dot represents the average signal from a single mouse, and error bars denote the standard deviation (N = 3 mice). **(C)** Mean (± SEM) RdLight1 photoactivation signal in the VLS in response to a pair of 4 ms blue light pulses (10 mW) separated by the indicated inter-pulse intervals. Mean paired pulse ratios are displayed (magnitude of pulse #2/pulse #1), with each dot representing a single mouse, and the error bars denoting the standard deviation (N = 3 mice).

## References

Akerboom J, Carreras Calderon N, Tian L, Wabnig S, Prigge M, Tolo J, Gordus A, Orger MB, et al. (2013). Genetically encoded calcium indicators for multi-color neural activity imaging and combination with optogenetics. Front Mol Neurosci 6: 2.

Berke JD (2018). What does dopamine mean? Nat Neurosci 21(6): 787–793.

Cachope R, Mateo Y, Mathur BN, Irving J, Wang HL, Morales M, Lovinger DM and Cheer JF (2012). Selective activation of cholinergic interneurons enhances accumbal phasic dopamine release: setting the tone for reward processing. Cell Rep 2(1): 33–41.

Chantranupong L, Beron CC, Zimmer JA, Wen MJ, Wang W and Sabatini BL (2023). Dopamine and glutamate regulate striatal acetylcholine in decision-making. Nature 621(7979): 577–585.

Da Silva JA, Tecuapetla F, Paixao V and Costa RM (2018). Dopamine neuron activity before action initiation gates and invigorates future movements. Nature 554(7691): 244–248.

Dana H, Mohar B, Sun Y, Narayan S, Gordus A, Hasseman JP, Tsegaye G, Holt GT, et al. (2016). Sensitive red protein calcium indicators for imaging neural activity. Elife 5.

Howe M, Ridouh I, Allegra Mascaro AL, Larios A, Azcorra M and Dombeck DA (2019). Coordination of rapid cholinergic and dopaminergic signaling in striatum during spontaneous movement. Elife 8.

Howe MW and Dombeck DA (2016). Rapid signalling in distinct dopaminergic axons during locomotion and reward. Nature 535(7613): 505–510.

Klapoetke NC, Murata Y, Kim SS, Pulver SR, Birdsey-Benson A, Cho YK, Morimoto TK, Chuong AS, et al. (2014). Independent optical excitation of distinct neural populations. Nat Methods 11(3): 338–346.

Kramer PF, Brill-Weil SG, Cummins AC, Zhang R, Camacho-Hernandez GA, Newman AH, Eldridge MaG, Averbeck BB and Khaliq ZM (2022). Synaptic-like axo-axonal transmission from striatal cholinergic interneurons onto dopaminergic fibers. Neuron 110(18): 2949–2960 e2944.

Krok AC, Maltese M, Mistry P, Miao X, Li Y and Tritsch NX (2023). Intrinsic dopamine and acetylcholine dynamics in the striatum of mice. Nature.

Liu C, Cai X, Ritzau-Jost A, Kramer PF, Li Y, Khaliq ZM, Hallermann S and Kaeser PS (2022). An action potential initiation mechanism in distal axons for the control of dopamine release. Science 375(6587): 1378–1385.

Lovinger DM, Mateo Y, Johnson KA, Engi SA, Antonazzo M and Cheer JF (2022). Local modulation by presynaptic receptors controls neuronal communication and behaviour. Nat Rev Neurosci 23(4): 191–203.

Mamaligas AA, Cai Y and Ford CP (2016). Nicotinic and opioid receptor regulation of striatal dopamine D2-receptor mediated transmission. Sci Rep 6: 37834.

Markowitz JE, Gillis WF, Jay M, Wood J, Harris RW, Cieszkowski R, Scott R, Brann D, et al. (2023). Spontaneous behaviour is structured by reinforcement without explicit reward. Nature 614(7946): 108–117.

Marshel JH, Kim YS, Machado TA, Quirin S, Benson B, Kadmon J, Raja C, Chibukhchyan A, et al. (2019). Cortical layer-specific critical dynamics triggering perception. Science 365(6453).

Matityahu L, Gilin N, Sarpong GA, Atamna Y, Tiroshi L, Tritsch NX, Wickens JR and Goldberg JA (2023). Acetylcholine waves and dopamine release in the striatum. Nat Commun 14(1): 6852.

Mohebi A, Collins VL and Berke JD (2023). Accumbens cholinergic interneurons dynamically promote dopamine release and enable motivation. Elife 12.

Mohebi A, Pettibone JR, Hamid AA, Wong JT, Vinson LT, Patriarchi T, Tian L, Kennedy RT and Berke JD (2019). Dissociable dopamine dynamics for learning and motivation. Nature 570(7759): 65–70.

Patriarchi T, Cho JR, Merten K, Howe MW, Marley A, Xiong WH, Folk RW, Broussard GJ, et al. (2018). Ultrafast neuronal imaging of dopamine dynamics with designed genetically encoded sensors. Science 360(6396).

Patriarchi T, Mohebi A, Sun J, Marley A, Liang R, Dong C, Puhger K, Mizuno GO, et al. (2020). An expanded palette of dopamine sensors for multiplex imaging in vivo. Nat Methods 17(11): 1147–1155.

Shaner NC, Lin MZ, Mckeown MR, Steinbach PA, Hazelwood KL, Davidson MW and Tsien RY (2008). Improving the photostability of bright monomeric orange and red fluorescent proteins. Nat Methods 5(6): 545–551.

Sulzer D, Cragg SJ and Rice ME (2016). Striatal dopamine neurotransmission: regulation of release and uptake. Basal Ganglia 6(3): 123–148.

Threlfell S, Lalic T, Platt NJ, Jennings KA, Deisseroth K and Cragg SJ (2012). Striatal dopamine release is triggered by synchronized activity in cholinergic interneurons. Neuron 75(1): 58–64.

Wang L, Shang S, Kang X, Teng S, Zhu F, Liu B, Wu Q, Li M, et al. (2014). Modulation of dopamine release in the striatum by physiologically relevant levels of nicotine. Nat Commun 5: 3925.

Wei W, Mohebi A and Berke JD (2022). A spectrum of time horizons for dopamine signals. BioRxiv.

Zhuo Y, Luo B, Yi X, Dong H, Miao X, Wan J, Williams JT, Campbell MG, et al. (2023). Improved green and red GRAB sensors for monitoring dopaminergic activity in vivo. Nat Methods.

